# Tristetraprolin prevents gastric metaplasia in mice by suppressing pathogenic inflammation

**DOI:** 10.1101/2021.01.21.427435

**Authors:** Jonathan T. Busada, Stuti Kadka, Kylie N. Peterson, Deborah J. Stumpo, Lecong Zhou, John A. Cidlowski, Perry J. Blackshear

## Abstract

Aberrant immune activation is associated with numerous inflammatory and autoimmune diseases and contributes to cancer development and progression. Within the stomach, inflammation drives a well-established sequence from gastritis to metaplasia, eventually resulting in adenocarcinoma. Unfortunately, the processes that regulate gastric inflammation and prevent carcinogenesis remain unknown. Tristetraprolin (TTP) is an RNA-binding protein that promotes the turnover of numerous pro-inflammatory and oncogenic mRNAs. Here, we utilized a TTP-overexpressing model, the TTPΔARE mouse, to examine whether TTP can protect the stomach from adrenalectomy (ADX)-induced gastric inflammation and spasmolytic polypeptide-expressing metaplasia (SPEM). We found that TTPΔARE mice were completely protected from ADX-induced gastric inflammation and SPEM. RNA sequencing revealed that TTP overexpression suppressed the expression of genes associated with the innate immune response. Finally, we show that protection from gastric inflammation was only partially due to suppression of *Tnf*, a well-known TTP target. Our results demonstrate that TTP exerts broad anti-inflammatory effects in the stomach and suggest that therapies that increase TTP expression may be effective treatments of pro-neoplastic gastric inflammation.

## Introduction

Gastric adenocarcinoma is the third leading cause of cancer deaths worldwide (1). Chronic inflammation is strongly correlated with gastric cancer development and is typically initiated by *Helicobacter pylori* infection or autoimmune gastritis (2, 3). Chronic inflammation causes atrophic gastritis and loss of the acid-secreting parietal cells (oxyntic atrophy), leading to the development of spasmolytic polypeptide-expressing metaplasia (SPEM) (4). These lesions are postulated to be the precursors of gastric adenocarcinoma (5, 6). Excessive expression of numerous proinflammatory cytokines, including IFNG, TNF, and IL1B, have been linked to SPEM development and gastric carcinogenesis (4, 7-9). However, the mechanisms that regulate gastric inflammation are poorly defined.

Tristetraprolin (TTP) is a member of a small family of RNA binding proteins and is encoded by the gene *Zfp36* (10, 11). Proteins of the TTP family are characterized by highly conserved tandem zinc finger domains which bind to AU rich-elements (AREs) in the 3’ UTR of target mRNAs (12, 13). TTP binding initiates de-adenylation and degradation of the target transcript (13). The ideal binding sequence, UUAUUUAUU or its variants, recognized by TTP, is found in a host of transcripts that encode pro-inflammatory cytokines, chemokines and oncogenes (14, 15). Loss of TTP expression increases the half-life of target mRNAs, and TTP expression is reduced or lost in numerous cancers, including gastric cancer (15, 16). Germline *Zfp36* KO mice exhibit numerous systemic inflammatory pathologies such as dermatitis, arthritis, autoimmunity, and myeloid hyperplasia, all of which are linked to aberrant expression of the proinflammatory cytokine TNF (17).

TTP expression is rapidly induced during inflammation, in which it regulates the intensity and duration of the inflammatory response (18). However, TTP expression is usually transient, in part due to binding sites within the TTP transcript that allow the TTP protein to negatively regulate its own expression (19). Mice with a germ-line deletion of a 136 base AU-rich element (ARE) within the TTP mRNA 3’UTR (TTPΔARE) have enhanced TTP mRNA stability and moderately increased TTP protein expression in their tissues, but are phenotypically normal during normal vivarium conditions (20). However, TTPΔARE mice have been shown to be resistant to experimental models of imiquimod-induced dermatitis, collagen antibody-induced arthritis, experimental autoimmune encephalomyelitis, bacterial gingivitis and dental bone erosion, inflammatory lung damage, experimental autoimmune uveitis, and chemically induced skin carcinogenesis (21-24).

We have previously demonstrated that glucocorticoids are master regulators of gastric inflammation. Systemic removal of endogenous glucocorticoids by ADX triggers massive, spontaneous gastric inflammation and SPEM (25). Adrenalectomy is a useful model to study factors that participate in gastric inflammation and SPEM development. In this study, we utilized the ADX model to investigate the effects of enhanced TTP expression on gastric inflammation and metaplasia. We found that increased, regulated, “whole body” TTP expression completely blocked the development of gastric inflammation and metaplasia after bilateral ADX. Surprisingly, protection from ADX-induced gastric inflammation was not recapitulated in *Tnf* KO mice, suggesting that TTP regulation of gastric inflammation is more complex than the suppression of a single proinflammatory cytokine. Our results suggest that treatments that increase TTP protein expression may effectively treat gastric inflammation and potentially protect against neoplasia development.

## Results

### TTP suppresses adrenalectomy-induced gastric inflammation

Gastric inflammation is associated with the development of gastritis, oxyntic atrophy, and metaplasia. TTP enhances the turnover of numerous proinflammatory mRNAs such as that encoding TNF (14, 17). We hypothesized that enhanced systemic TTP expression could protect mice from gastric inflammation and metaplasia. To test this hypothesis, we utilized the TTPΔARE mice, in which a 136 base AU-rich instability region was deleted from the 3’ UTR of the gene encoding TTP, *Zfp36* (20). As previously reported in other tissues (20), we confirmed that germline deletion of the ARE region results in the accumulation of TTP mRNA in the mouse gastric fundus at 2 months of age (Figure 1A). We previously showed that adrenalectomy (ADX) rapidly induces spontaneous gastric inflammation and SPEM (25). We utilized bilateral ADX to assess gastric inflammation and SPEM development in TTPΔARE mice (Figure 1B). As expected, WT control mice exhibited prominent inflammation within the gastric corpus 2 months post-ADX (Figure 1C). In contrast, both TTPΔARE heterozygous and homozygous mice were protected from increased inflammation. In addition to stomach inflammation, ADX WT mice developed splenomegaly (Figure 1D), a classic feature of ADX in rodents (26). However, TTPΔARE heterozygous and homozygous spleen weights did not significantly differ from WT mice 2 months after ADX. These data indicate that increased systemic TTP expression from normally regulated *Zfp36* can protect the stomach from the chronic inflammation caused by ADX.

**Figure 1.**
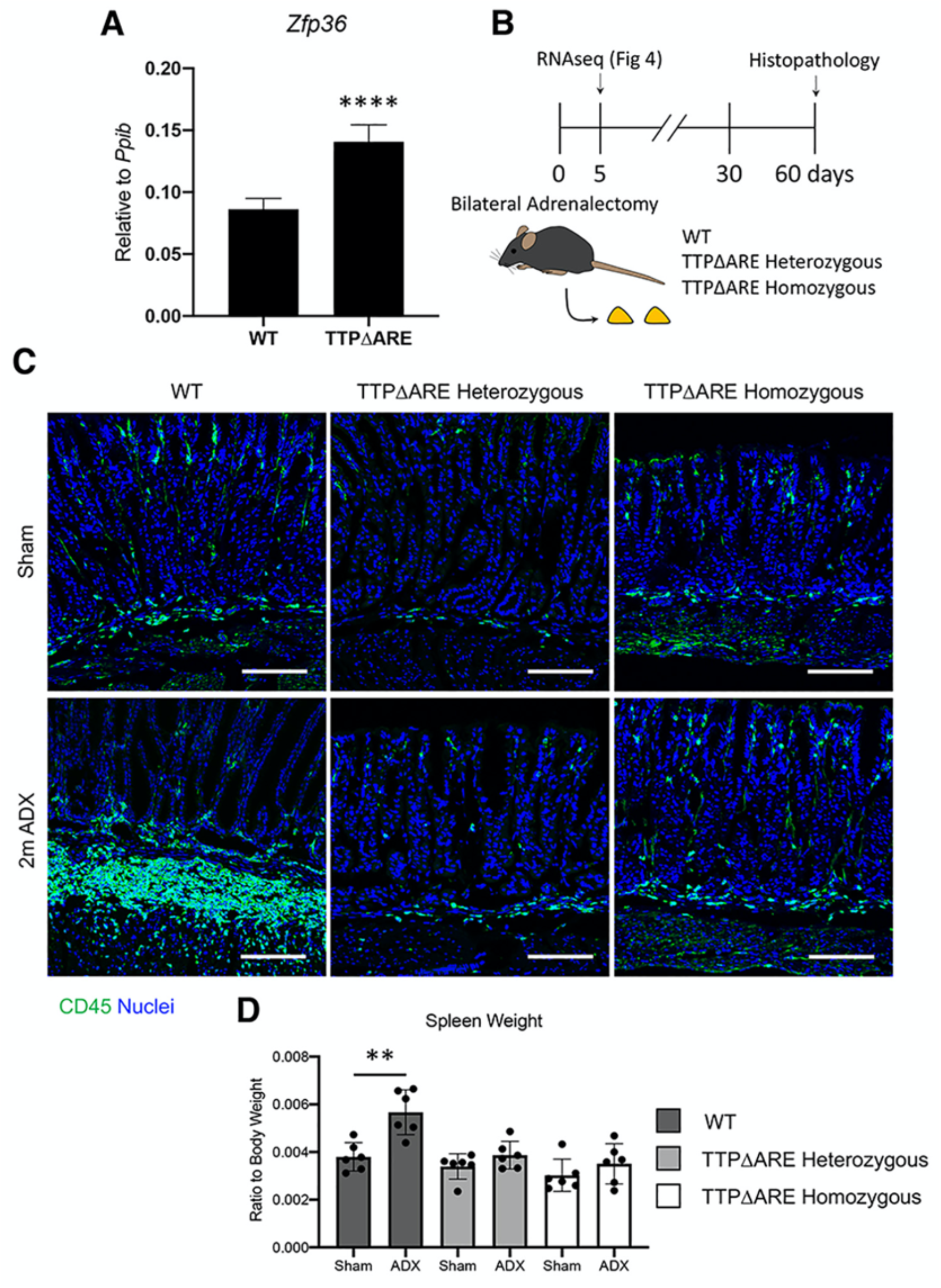
Elevated levels of TTP prevent adrenalectomy-induced gastric inflammation. (**A**) qRT-PCR for TTP mRNA within the gastric corpus. (**B**) Experimental model. (**C**) Representative immunostaining of the gastric corpus from WT, TTP ΔARE heterozygous, and TTP ΔARE homozygous mice euthanized 2 months after sham surgery or adrenalectomy. Gastric sections were stained with CD45 antibodies and nuclei were stained with DAPI. Scale bars are 100 µm; *n*≥7 mice/group. (A and D) Data are mean ± SD; *P*-values were determined by unpaired student T-test (A) or one-way ANOVA with post hoc Tukey’s t-test (D). ** *P*≤0.01; \*\*\*\**P*≤0.0001.

### TTP protects mice from spasmolytic polypeptide-expressing metaplasia development

SPEM develops in response to glandular damage such as oxyntic atrophy and is a putative precursor of gastric adenocarcinoma (27). Inflammation potently induces SPEM development (25). Because TTPΔARE mice were resistant to ADX-induced inflammation, we asked whether increased TTP expression could prevent the development of oxyntic atrophy and metaplasia. The gross morphology of sham-operated TTPΔARE heterozygous and homozygous mice was indistinguishable from sham-operated WT mice (Figure 2A), and there were no significant differences in the number of parietal cells or chief cells (Figure 2B). Two months after ADX, WT mice had lost 82% of their parietal cell population and 99% of their mature chief cells (Figure 2). Moreover, WT mice exhibited prominent mucous cell hyperplasia within the gastric corpus, identified by an increase in *Griffonia simplicifolia* (GSII) lectin staining, which binds to mucin 6 (MUC6). In contrast to ADX-WT mice, neither TTPΔARE heterozygous nor homozygous mice exhibited a significant change in their parietal and chief cell populations, and both genotypes had normal gastric morphology 2 months after ADX (Figure 2).

**Figure 2.**
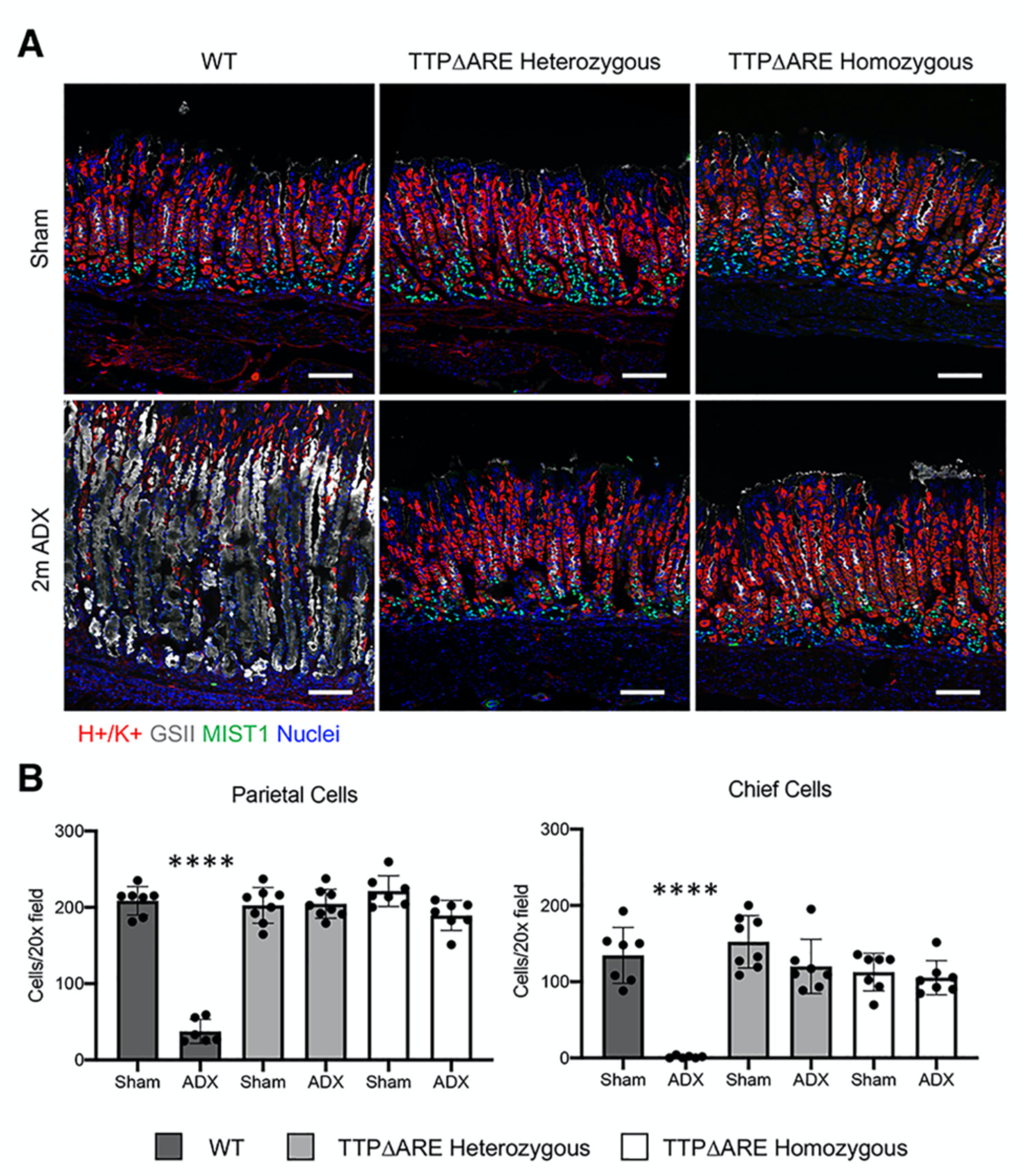
Elevated levels of TTP protect the stomach from adrenalectomy-induced pathogenic inflammation. **(A)** Representative immunostaining of the gastric corpus from WT, TTPΔARE heterozygous, and TTPΔARE homozygous mice euthanized 2 months after sham surgery or adrenalectomy. Gastric sections were probed for ATP4B (parietal cells, red), MIST1 (chief cells, green), and GSII lectin (mucous neck cells, grey). Nuclei were stained with DAPI. Scale bars are 100 µm. (**B**) Quantitation of the number of parietal cells and chief cells observed per 20x field (*n*≥6 mice/group). Data are mean ± SD; *P*-values were determined by one-way ANOVA with post hoc Tukey’s t-test. \*\*\*\**P*≤0.0001.

Oxyntic atrophy, loss of the mature chief cell marker MIST1, and expansion of GSII+ cells are defining characteristics of SPEM. We confirmed SPEM development by immunostaining for the *de novo* SPEM marker CD44v9, a splice variant of CD44 that is induced during SPEM (28). While there was widespread staining of CD44v9 in ADX WT mice, CD44v9 was not detected within the gastric glands of ADX TTPΔARE mice (Figure 3A). In addition, we performed qRT-PCR on a panel of transcripts from the advanced SPEM-associated genes, *Cftr, Wfdc2*, and *Olfm4* (29, 30). Consistent with the increase in CD44v9 staining, there was significant induction of all three advanced SPEM markers in ADX WT mice (Figure 3B). However, these transcripts did not significantly increase in TTPΔARE homozygous mice. These results demonstrate that increased TTP expression protected the mice from oxyntic atrophy and SPEM development.

**Figure 3.**
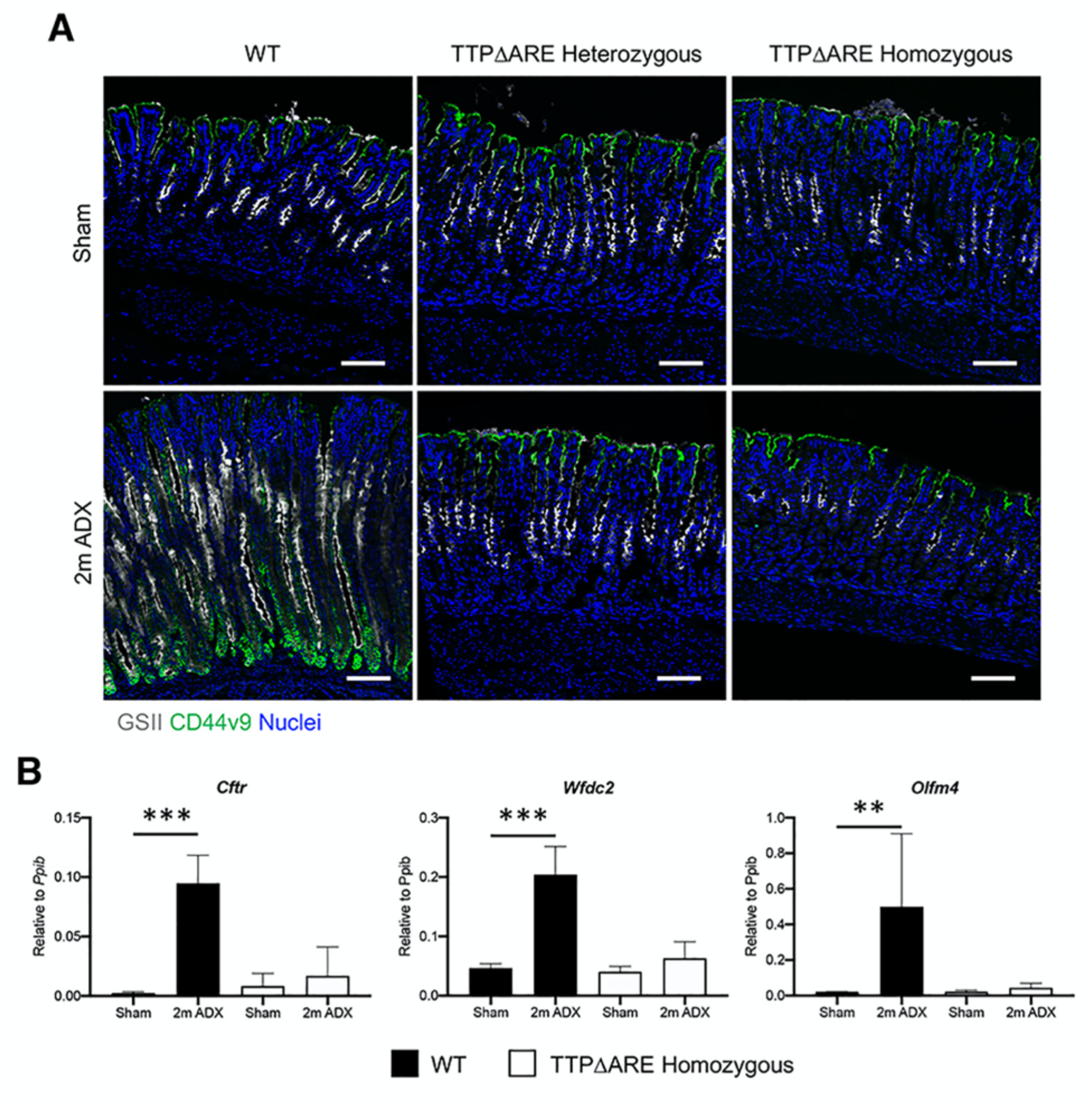
Elevated levels of TTP prevent SPEM development. **(A)** Representative immunostaining of the gastric corpus from WT, TTPΔARE heterozygous, and TTPΔARE homozygous mice euthanized 2 months after sham surgery or adrenalectomy. Gastric sections were probed for the SPEM marker CD44v9 (green) and the lectin GSII (mucous neck cells, grey). Nuclei are stained with DAPI. Scale bars are 100 µm, *n*≥6 mice/group. (**B**) Quantitative RT-PCR of the indicated SPEM marker genes using RNA isolated from the gastric corpus (*n*≥4 mice/group). Data are mean ± SD; *P*-values were determined by one-way ANOVA with post hoc Tukey’s t-test. * *P*≤0.05, \*\**P*≤0.001, \*\*\**P*≤0.001 \*\*\*\**P*≤0.0001.

### TTP suppresses the induction of proinflammatory gene mRNAs after adrenalectomy

Because TTPΔARE mice were protected from ADX-induced gastric inflammation and SPEM, we next utilized RNAseq to examine their gastric transcriptomes 5 days after ADX (Figure 1B). We used this early time after ADX to avoid secondary changes due to the anatomical alterations seen in longer term ADX mice. RNAseq revealed significant increases in inflammatory gene expression 5 days after ADX in WT mice. Gene set enrichment analysis (GSEA) comparing the sham WT vs. ADX WT groups revealed significant enrichment of mRNAs associated with the “activation of the inflammatory response” gene ontology pathway (Figure 4A). Surprisingly, there was also significant enrichment of inflammatory genes 5 days after ADX in TTPΔARE homozygous mice. However, the normalized enrichment score (NES) was 6.36 in the WT group compared to 5.02 in the TTPΔARE group, suggesting moderately increased inflammation within the WT group. Moreover, a comparison of the ADX WT group to the ADX TTPΔARE group demonstrated greater activation of inflammatory response pathways in ADX WT mice (Figure 4A). Next, we ranked the GSEA data and found that the “GO innate immune response” pathway was the 7^th^ highest activated pathway in the WT group (NES=5.32, Figure 4B). In contrast, this pathway was ranked 46^th^ in the TTPΔARE group (NES=3.97). Comparison of the WT ADX group to the TTPΔARE ADX group demonstrated significant positive enrichment (Figure 4B), suggesting increased innate immune system activation in WT ADX mice.

**Figure 4.**
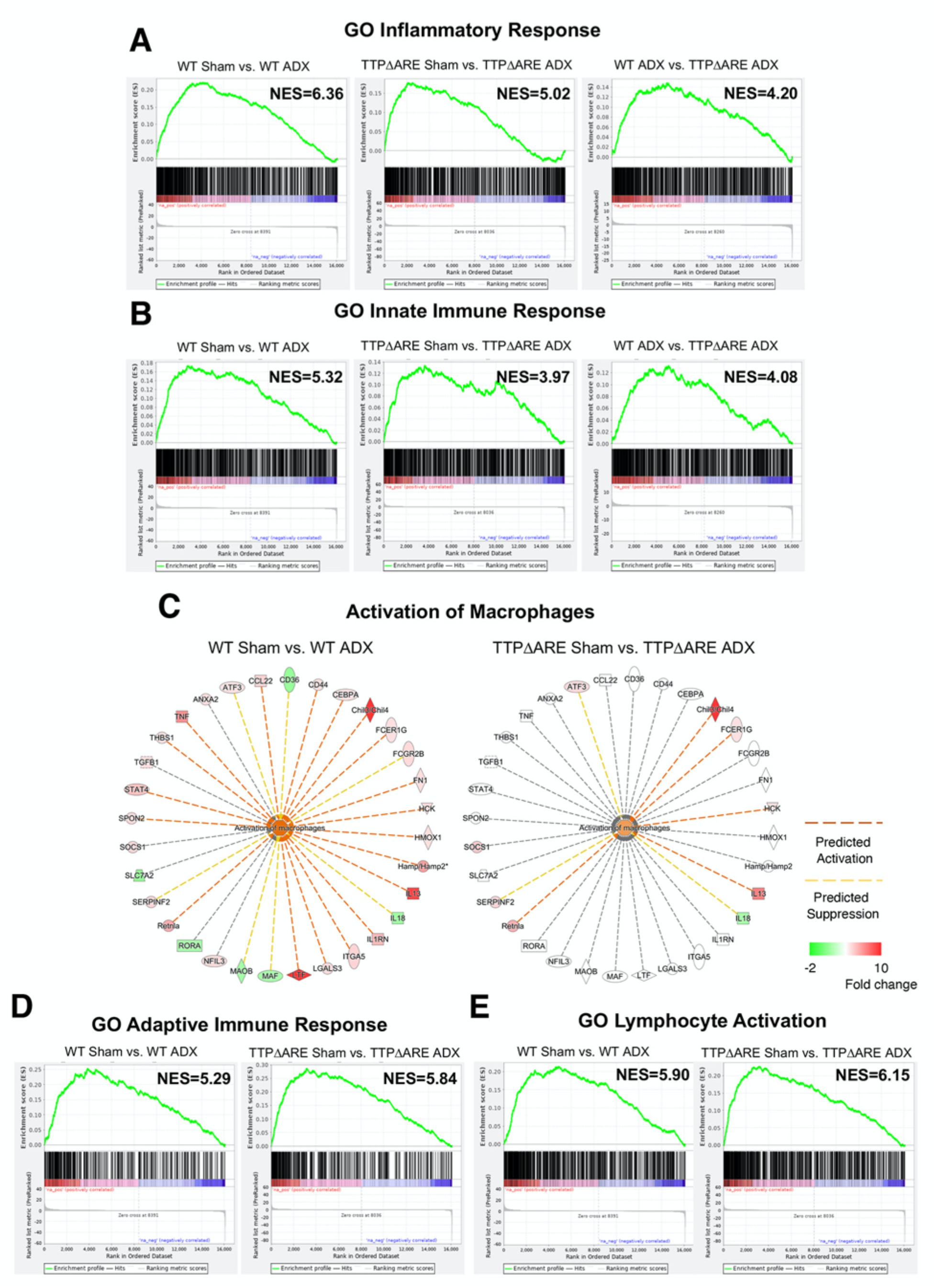
Increased TTP expression elicits broad suppression of inflammatory genes within the stomach of adrenalectomized mice. Shown are RNA-Seq data from total cellular RNA isolated from the gastric corpus of WT and TTPΔARE homozygous mice euthanized 5 days after sham surgery or adrenalectomy. (**A-B** and **D-E**) Gene Set Enrichment Analysis (GSEA) of the significantly regulated genes comparing adrenalectomized WT mice to adrenalectomized TTP mice. (**C**) Ingenuity Pathway Analysis of the total number of significantly regulated genes in sham vs. adrenalectomized WT mice and TTP mice, respectively. n=4 mice/group.

Macrophages have previously been shown to be required to induce SPEM development (25, 31). Therefore, we next analyzed the differentially expressed gene (DEG) lists using Ingenuity Pathway Analysis (IPA) to assess transcripts associated with macrophage activation. IPA predicted significant activation of the “Activation of Macrophages” pathway in ADX WT mice (activation z-score=2.43; Figure 4C). However, this pathway was not significantly activated in ADX TTPΔARE mice. Importantly, GSEA demonstrated that pathways associated with adaptive immunity such as the “GO adaptive immune response” (Figure 4D) and “lymphocyte activation” (Figure 4E) were equivalently activated in both WT and TTPΔARE mice. These results are consistent with published reports that mature lymphocytes are dispensable for inducing SPEM development (25).

### Transcripts containing adenylate-uridylate-rich elements are only a small portion of the ADX-induced genes

TTP is an RNA binding protein that binds to adenylate-uridylate-rich target sequences in mRNAs before promoting the turnover of those mRNAs. RNAseq revealed 760 DEGs between the sham WT and ADX WT groups. In contrast, there were only 490 DEGs between the sham TTPΔARE mice and ADX TTPΔARE groups (Figure 5A). Of the DEGs, 189 genes were regulated in both groups. We screened the transcripts that were upregulated by ADX in the WT group for the presence of ideal TTP binding sequences (UAUUUAU and UAUUUUAU). We identified 94 mRNAs that contained a potential TTP binding motif (Figure 5B). Upregulation of 93 of these transcripts was significantly blunted in ADX TTPΔARE mice, indicating that TTP may enhance the degradation of these transcripts. Importantly, among the 94 ARE-containing transcripts, there were established TTP targets such as the mRNA encoding *Tnf*, as well as numerous other inflammatory genes, including *Il13*, which is required for SPEM development (32). These data suggest that TTP directly regulates approximately 33% of DEG in the stomachs of ADX WT mice. However, there are likely also numerous genes that are indirectly regulated by TTP.

**Figure 5.**
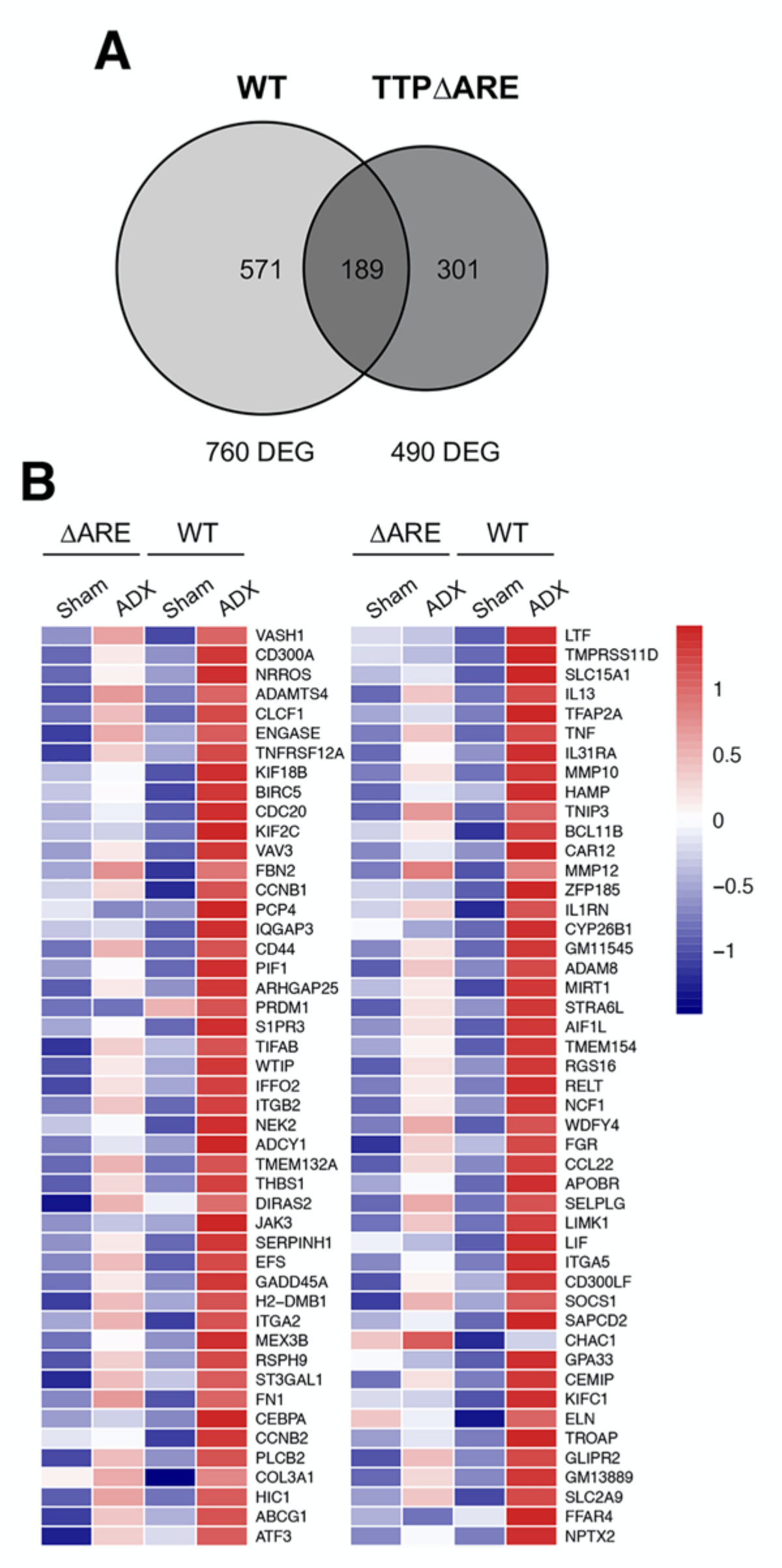
ARE-containing mRNAs are a small proportion of the differentially expressed genes in the stomach. **(A)** Venn diagram of the total number of DEGs in the gastric corpus 5 days after ADX. (**B**) Heatmap visualizing AU-rich element-containing mRNAs in the indicated data-sets. Transcripts were significantly upregulated in ADX WT mice compared to sham WT mice. Scale is z-score min/max within the dataset.

### Tnf knockout mice are partially protected from ADX-induced SPEM

Tumor necrosis factor alpha (TNF) is a prominent proinflammatory cytokine that is produced by macrophages and other leukocytes. Aberrant TNF production is associated with inflammatory disease within the GI tract, and may increase the risk of gastric cancer (9, 33). Moreover, *Tnf* mRNA is an established TTP target, and germline *Zfp36* KO mice have systemic inflammation attributed in part to excessive TNF (12, 17). We hypothesized that suppression of *Tnf* in TTPΔARE mice may protect against ADX-induced inflammation and metaplasia. Therefore, we adrenalectomized *Tnf* KO mice and assessed their stomachs 2 months after surgery. Interestingly, *Tnf* KO mice exhibited only intermediate protection from SPEM (Figure 6A-B). In ADX *Tnf* KO mice, there were regions of the gastric corpus that appeared identical to sham controls, with the normal complement of parietal and chief cells, and that were negative for the SPEM marker CD44v9 (Figure 6A). In contrast, other regions of the lesser curvature appeared identical to sections from the ADX WT mice (Figure 6A, far right panel). We quantitated the number of parietal and chief cells present in both “normal” and “SPEM” regions. Quantitation revealed that while ADX *Tnf* KO mice exhibited a significant loss of parietal and chief cells relative to sham controls, these effects were significantly diminished compared to ADX WT mice (Figure 6B). Surprisingly, *Tnf* KO completely rescued the splenomegaly observed in ADX WT mice (Figure 6C-D). Taken together, these data indicate that while *Tnf* contributes to SPEM development, there are likely redundant mechanisms that control pathogenic gastric inflammation. Moreover, these results demonstrate that TTP’s protective effects in the stomach are the result of broad anti-inflammatory effects beyond the simple suppression of *Tnf*.

**Figure 6.**
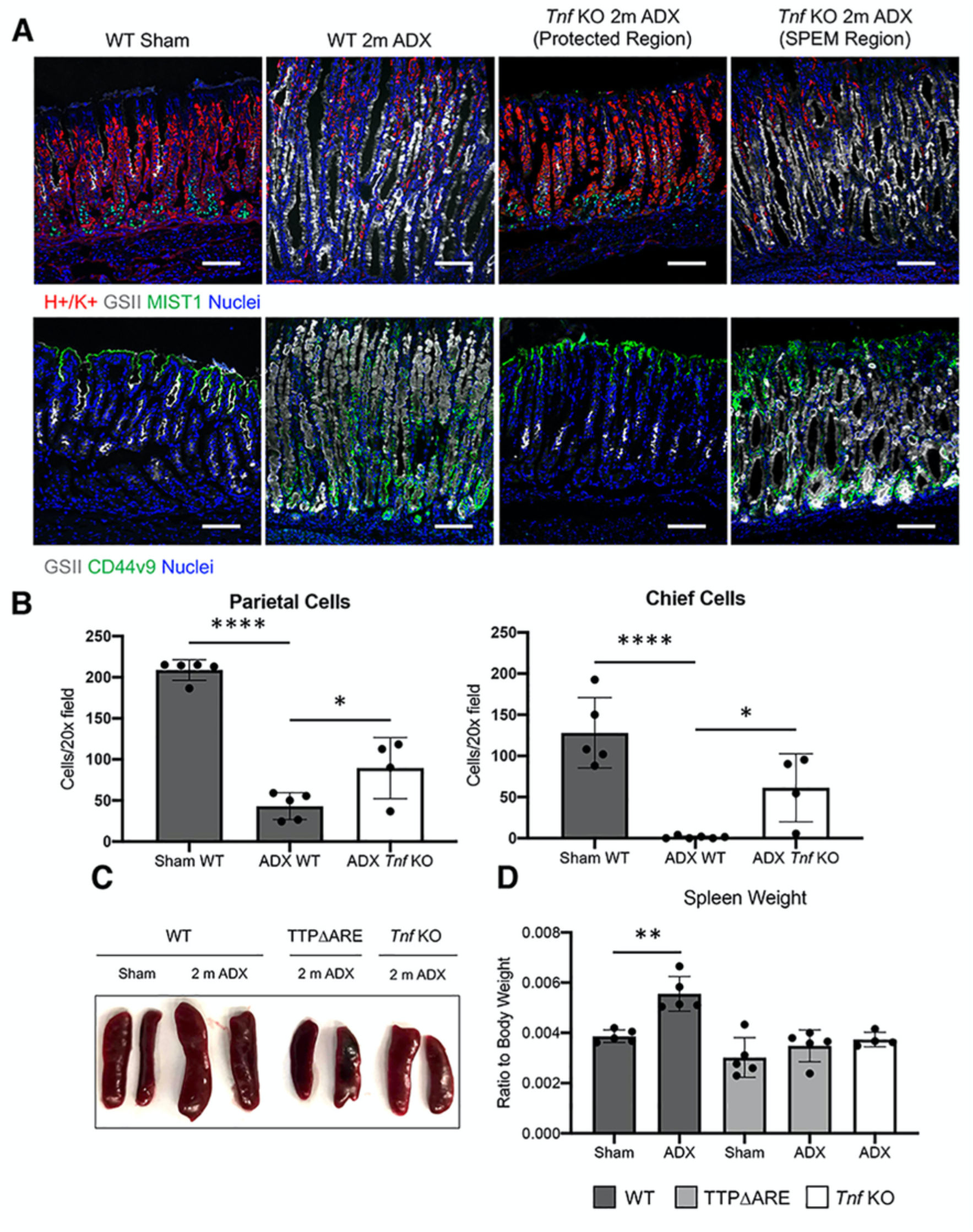
*Tnf* KO mice are partially protected from adrenalectomy-induced gastric inflammation. (**A**) Representative images of stomachs taken from WT and *Tnf* KO mice euthanized 2 months after sham surgery or adrenalectomy. Scale bars are 100 µm. (**B**) Quantitation of the number of parietal cells and chief cells observed per 20x field (*n*≥4 mice/group). (**C**) Representative images of spleens from mice euthanized 2 months after sham surgery or ADX (**D**) Ratio of spleen weight normalized to total body weight. Data are mean ± SD; *P*-values were determined by one-way ANOVA with post hoc Tukey’s t-test. * *P*≤0.05, \*\**P*≤0.001, \*\*\**P*≤0.0001.

## Discussion

Post-transcriptional regulation of gene expression by RNA binding proteins is critical for maintaining cellular and tissue homeostasis. Dysregulation of RNA binding proteins is associated with a host of diseases including cancer (34). *Zfp36* encodes a zinc finger RNA binding protein, TTP, that binds to ARE-containing mRNAs and destabilizes them by recruiting deadenylases, thus promoting mRNA decay (35). It has been estimated that about 26% of human mRNA 3’UTRs contain at least a single minimal TTP family binding site, UAUUUAU or UAUUUUAU (36), and disruption of TTP family members has been associated with inflammatory disorders and cancer (15, 37-39). TTP is a critical regulator of numerous proinflammatory cytokines. TTP KO mice develop multi-system inflammatory disease that is largely due to excessive TNF expression (15, 17, 40). In contrast, increased TTP expression confers resistance to numerous inflammatory pathologies including arthritis and dermatitis (20-22). Here, we report that knockin mice that have regulated increases in TTP levels throughout the body are protected from ADX-induced gastric inflammation and SPEM. Our results suggest that TTP could be a master regulator of gastric inflammation, and therapies that lead to increased TTP protein levels may be effective at treating gastric inflammation.

Chronic inflammation is strongly associated with gastric cancer development. Within the stomach, inflammation induces a well-defined histopathological progression in which stomach damage leads to gastric atrophy, metaplasia, dysplasia, and adenocarcinoma (6). SPEM is a potentially preneoplastic form of metaplasia that develops in response to damage within the gastric corpus that may also serve as a healing mechanism (5, 27). However, in the setting of prolonged damage, such as during chronic inflammation, SPEM becomes increasingly proliferative and may eventually progress towards carcinogenesis. We found that TTP overexpression protected mice from gastric inflammation and SPEM development. We utilized ADX as a model to challenge the TTPΔARE mice. In WT mice, ADX triggered massive spontaneous inflammation of the gastric corpus followed by SPEM development. Both homozygous and heterozygous TTPΔARE mice were completely protected from ADX-induced gross inflammation and SPEM development. We previously reported that suppressing gastric inflammation by depleting macrophages in ADX WT mice protects them from SPEM development (25). Thus, it is likely that TTP prevents SPEM development by suppressing inflammation. Our results suggest that therapies that elicit even a modest increase in TTP expression may be effective in controlling gastric inflammation.

Several studies have reported that macrophages are critical for inducing SPEM development (9, 25, 31). TTP primarily functions by binding to specific AU-rich elements within the 3’ UTR of target mRNAs, eventually promoting the degradation of the mRNA (41). Our RNAseq studies demonstrated that TTP potently suppressed genes associated with macrophage activation in ADX mice. Importantly, TTP regulates the expression of IL33, IL13, and TNFα, cytokines that have been implicated in inducing SPEM development (9, 16, 32). Our RNAseq data revealed that 33% of DEG transcripts in ADX WT mice contained potential TTP binding sites. Among these transcripts were several that encoded proinflammatory cytokines such as TNF and IL13 (9, 32). However, given that TTP can regulate other cellular pathways, including those involving NFκB (42, 43), it is likely that TTP can indirectly regulate the expression of additional inflammatory genes within the stomach.

Surprisingly, despite the almost complete suppression of inflammatory infiltrates into the stomachs of ADX TTPΔARE mice, we found striking upregulation of numerous inflammatory transcripts and pathways. Increased TTP was effective in suppressing the innate immune response, while pathways associated with the adaptive immune response were not significantly affected. It has been postulated that TTP preferentially regulates the innate immune response. However, while myeloid-specific TTP KO mice have an abnormal inflammatory response when challenged with lipopolysaccharide, they do not phenocopy the spontaneous inflammatory pathologies that develop in the whole body TTP KO (44). Several studies have found that lymphocytes are dispensable for inducing SPEM development (9, 25, 31). Thus, even if TTP primarily suppresses the innate immune system it may be inconsequential for SPEM development.

Aberrant TNF production is associated with numerous inflammatory pathologies of the gastrointestinal tract (45). *Helicobacter pylori* infection potently induces TNF production, and *Tnf* KO mice are protected from SPEM development in some mouse models (9, 33). Thus, TNF may contribute to gastric carcinogenesis. *Tnf* mRNA is a well-known TTP target, and the numerous inflammatory pathologies that develop in TTP KO mice were rescued by treatment with TNF neutralizing antibodies (17, 41) or by breeding to TNF receptor deficient mice (40). While macrophages produce large amounts of TNF and are critical for driving SPEM development (44), we hypothesized that TTP suppression of *Tnf* was the underlying mechanism by which TTPΔARE mice were protected from ADX-induced gastric inflammation. Surprisingly, we found that *Tnf* KO mice were at least partially susceptible to ADX-induced gastric inflammation and metaplasia. Interestingly, *Tnf* KO mice did not develop ADX-induced splenomegaly. These results demonstrate tissue-specific roles for TTP in regulating inflammation, and suggest that TTP’s anti-inflammatory role in the stomach is more complex than the suppression of a single proinflammatory cytokine.

Regulation of inflammation is multifaceted, occurring at the transcriptional level, post-transcriptional level, and beyond. We have previously demonstrated that glucocorticoids are critical transcriptional regulators of gastric inflammation (25). Here, we report that increased expression of the RNA binding protein TTP protects mice from gastric inflammation and metaplasia. Our results suggest that TTP is a master regulator of the gastric inflammatory response and may be a useful therapeutic target for treating gastric inflammatory disease. Recent reports have found that TTP expression is decreased in gastric cancer samples (16). Thus, there is a need for continued study into the role of TTP in suppressing gastric inflammation and carcinogenesis.

## Materials and Methods

### Animal care and treatment

All mouse studies were performed with approval by the NIEHS animal care and use committee. C57BL/6J (#000664) mice were purchased from the Jackson Laboratories (Bar Harbor, MA). TTPΔARE mice were generated as previously described and were maintained on a congenic C57Bl/6 genetic background (20). Mice were administered standard chow and water *ab libitum* and maintained in a temperature and humidity-controlled room with standard 12-hour light/dark cycles. Sham, adrenalectomy, and castration surgeries were performed at 8 weeks of age by the NIEHS Comparative Medicine Branch. Following adrenalectomy, mice were maintained on 0.85% saline drinking water to maintain ionic homeostasis.

### Histology

Mice were euthanized by cervical dislocation at the indicated timepoints. Stomachs were removed and opened along the greater curvature and washed in phosphate buffered saline to remove gastric contents. Stomachs were fixed overnight in 4% paraformaldehyde at 4°C and then cryopreserved in 30% sucrose and embedded in optimal cutting temperature (OCT) media. Histology and cell quantitation was performed as previously described (25). Briefly, 5 µm stomach cryosections were incubated with antibodies against the H+/K+ ATPase α subunit (Clone 1H9, MBL International Corporation, Woburn, MA), MIST1 (clone D7N4B, Cell Signaling Technologies, Danvers, MA), CD45 (clone 104, Bioledgend, San Diego, CA), or CD44v9 (Cosmo Bio, Tokyo, Japan) for 1 hour at room temperature or overnight at 4°C. After washing in PBS with 0.1% Triton X 100, sections were incubated in secondary antibodies for 1 hour at room temperature. Fluorescent conjugated *Griffonia simplicifolia* lectin (GSII; Thermo Fisher Scientific, Waltham, MA) was added with secondary antibodies. Sections were mounted with Vectastain mounting media containing DAPI to visualize nuclei (Vector Laboratories Burlingame, CA). Images were obtained using a Zeiss 710 confocal laser-scanning microscope equipped with Airyscan (Carl-Zeiss GmbH, Jena, Germany) and running Zen Black imaging software. Parietal cells and chief cells were quantitated as previously described (25) using confocal micrographs captured using a 20x microscope objective and 1 µm thick optical sections. Cells were counted using the ImageJ (National Institutes of Health, Bethesda, MD) count tool. Cells that stained positive with anti-H+/K+ antibodies were identified as parietal cells, while cells that stained positive with anti-MIST1 antibodies and were GSII negative were identified as mature chief cells. Counts were reported as the number of cells observed per 20x field. Images were chosen that contain gastric glands cut longitudinally.

### RNA isolation and qRT-PCR

RNA used for qRT-PCR and RNAseq was isolated from a 4mm biopsy of the gastric corpus lesser curvature. RNA was extracted in TRIzol (Thermo Fisher Scientific) and precipitated from the aqueous phase using 1.5 volumes of 100% ethanol. The mixture was transferred to a RNeasy column (Qiagen, Hilden, Germany), and the remaining steps were followed according to the RNeasy kit manufacturer’s recommendations. RNA was treated with RNase free DNase I (Qiagen) as part of the isolation procedure. Reverse transcription followed by qPCR was performed in the same reaction using the Universal Probes One-Step PCR kit (Bio-Rad Laboratories, Hercules, CA) and the Taqman primers *Cftr* (Mm00445197_m1), *Olfm4* (Mm01320260_m1), *Wfdc2* (Mm00509434_m1), and *Zfp36* (Mm00457144_m1) (Thermo Fisher Scientific) on a Quantstudio 6 (Thermo Fisher Scientific). Messenger RNA levels were normalized to the reference gene *Ppib* (Mm00478295_m1).

### RNA Sequencing

RNA was isolated 5 days post sham surgery or adrenalectomy as described above. Four mice were used for each experimental group. Indexed samples were sequenced using the 75 bp paired-end protocol via the NextSeq500 (Illumina) per the manufacturer’s protocol. Raw reads (27-41 million pairs of reads per sample) were filtered using a custom perl script and the cutadapt program (v2.8) to remove low quality reads and adapter sequences. Preprocessed reads were aligned to the UCSC mm10 reference genome using *STAR* (v2.7.0f) with default parameters (46). The quantification results from *featureCounts* (available in Subread software, v1.6.4) were then analyzed with the Bioconductor package DESeq2, which fits a negative binomial distribution to estimate technical and biological variability (47). Comparisons were made between sham WT vs. ADX WT, sham TTPΔARE vs. ADX TTPΔARE, and ADX WT vs. TTPΔARE. An abundance cutoff was used so that only transcripts were evaluated whose average expression in the WT samples was greater than 0.1 FPKM. A transcript was considered differentially expressed if the *adjusted p* -value was <0.05 and its expression changed ≤-1.5 or ≥1.5 fold. Lists of significant transcripts were further analyzed using Ingenuity Pathway Analysis (version 01-18-05, Qiagen). Enrichment or overlap was determined by IPA using Fisher’s exact test (p < 0.05). Gene set enrichment analysis (GSEA) was performed using GSEA v4.0.3 software and Molecular Signatures Database (MSigDB) v7.0 (48). Transcripts were pre-ranked based on p-value and their fold change of gene expression. This application scores a sorted list of transcripts with respect to their enrichment in selected functional categories (KEGG, Biocarta, Reactome and GO). The significance of the enrichment score was assessed using 1000 permutations. Benjamini and Hochberg’s false discovery rate (FDR) was calculated for multiple testing adjustments. q ≤ 0.05 were considered significant. The heatmap was generated with the mean expression values of the 94 selected genes. The expression values were log2 transformed before subjecting to heatmap generation with scale by row in the pheatmap function available in R package pheatmap. The RNA-seq data are available in the Gene Expression Omnibus repository at the National Center for Biotechnology Information (https://www.ncbi.nlm.nih.gov/geo/) (accession number: GSE164349).

### Statistical analysis

All error bars are ± the standard deviation of the mean. The sample size for each experiment is indicated in the figure legends. Experiments were repeated a minimum of two times. Statistical analyses were performed using one-way ANOVA with post-hoc Tukey’s t-test when comparing 3 or more groups or by unpaired t-test when comparing 2 groups. Statistical analysis was performed by Graphpad Prism 8 software. Statistical significance was set at p≤0.05. Specific p values are listed in the figure legends.

## Abbreviations

GR: glucocorticoid receptor
SPEM: spasmolytic polypeptide expressing metaplasia
TTP: Tristetraprolin
AREs: adenylate-uridylate rich elements
ADX: adrenalectomy

## Acknowledgments

This research was supported by a Postdoctoral Research Associate (PRAT) fellowship from the National Institute of General Medical Sciences 1Fi2GM123974 (J.T.B.), by West Virginia University start-up funds (J.T.B), and by the Intramural Research Program of the NIH/NIEHS 1ZIAES090057 (J.A.C) and 1ZIAES090080 (P.J.B.). Confocal microscopy was performed in the West Virginia University Microscope Imaging Facility which is supported by NIH grants P30GM103488, P20GM103434, and U54GM10942. The authors thank the NIEHS Comparative Medicine Branch, Epigenomes Core Laboratory, and Fluorescence Microscopy and Imaging Center for their assistance. We also thank Michael Fessler and Donald Cook for critical reading of the manuscript.

